# Proteolytic processing of LRP2 on RPE cells regulates BMP activity to control eye size and refractive error

**DOI:** 10.1101/365817

**Authors:** Ross F. Collery, Brian A. Link

## Abstract

Mutations in LRP2, a transmembrane receptor, cause ocular enlargement and high-myopia. LRP2 is expressed by the RPE and eye ciliary epithelia, binding many extracellular ligands, including Bmp4 and Shh. Signaling mediated by LRP2 is very context-dependent, and how multiple pathways are coordinated is unknown. Transcriptome analyses of ocular tissues revealed that controlled, sustained BMP signaling from the RPE is critical for normal eye growth and emmetropia (proper refraction). Using zebrafish, we demonstrate that BACE sheddase-dependent LRP2 cleavage produces a soluble domain that binds BMP4, inhibiting its signaling. We propose that controlled proteolytic cleavage of LRP2 makes two ligand-binding receptor forms available: a soluble BMP trap, and a membrane-bound RPE signaling facilitator. By modulating LRP2 cleavage, cells can fine-tune and coordinate multiple signaling pathways, as well as growth and turnover of the extracellular matrix, control of which is important to maintain proper eye size. This data supports the concept that LRP2 acts as a homeostasis node that buffers and integrates diverse signaling to regulate emmetropic eye growth.

**Author Summary:** For proper focusing and normal vision, the axial length of the eye needs to match the refractive power of the lens. This is achieved by fine-tuning multiple signaling pathways to regulate the shape of the eye primarily by remodeling of the sclera, the outermost layer of the eye. This process is termed emmetropization. Emmetropization cues are initiated by visual input, but how signals are transduced from the photoreceptors across the retinal pigment epithelium to the sclera is incompletely understood. Here we show that cleavage of Lrp2, a large receptor expressed on RPE cells in the eye, alters BMP signaling, which contributes to proper eye size control. Dysregulation of BMP signaling by a) absence of Lrp2 in mutant zebrafish or b) overexpression of BMP antagonists from the RPE both cause eye enlargement and myopia. Understanding how regulated cleavage of Lrp2 affects paracrine signaling provides critical insight to emmetropization, raising the possibility for development of therapeutic agents to combat the epidemic incidence of refractive error.

## Introduction

LRP2/megalin is a large (>520 kD) transmembrane protein of the LDL-receptor related protein family that is expressed in retinal pigment epithelium (RPE) and ciliary epithelium of the eye (1–3), as well as on other absorptive epithelial cells such as proximal tubules in the kidney (4,5). Mutations in human *LRP2* lead to Donnai-Barrow syndrome, characterized by orbital hypertelorism, buphthalmia, high-grade myopia, agenesis of the corpus callosum, deafness, low molecular weight proteinuria and diaphragmatic hernia (6–8). Lrp2 binds a large number of proteins via four LDL type A domains. In the eye, Lrp2 has recently been shown to bind Sonic Hedgehog (Shh), preventing interaction with Patched1, and act as a clearance receptor to inhibit Shh signaling (9). In the absence of Lrp2, the optic cup margin shows extended proliferation, resulting in ciliary body hyperplasia. In the forebrain, however, Lrp2 functions as an auxiliary receptor with Patched1 to de-repress Smoothened activity and facilitate Shh signaling (10). Loss of Lrp2 in the forebrain results in holoprosencephaly and phenocopies Shh mutants. Forebrain deletion of Lrp2 also results in augmented BMP2/4/7 signaling (11). In that study, in vitro assays demonstrated Lrp2 can bind BMP4 and facilitate lysosomal degradation of the ligand. It is therefore clear that Lrp2 impacts signaling and homeostasis in context-dependent ways. As further example of the complexity of its regulation and function, LRP2 is cleaved in cultured kidney-derived cells by regulated intramembrane proteolysis (RIP), where the intracellular C-terminal domain can enter the nucleus and regulate gene expression (12,13). However, in vivo, over-expression of the C-terminus of Lrp2 in renal proximal tubules did not alter global gene expression nor result in obvious phenotype changes (14). Potentially, then, the extracellular domain may be the relevant bioactive fragment upon proteolytic processing. Indeed, soluble forms of LRP2 corresponding to extracellular fragments have also been reported, and are likely generated through cleavage at an extracellular site (15–19). Though the extracellular Lrp2-processing enzymes have not been identified, proteolysis has been predicted to be regulated by a sheddase (12). Since LRP2-mediated endocytosis is only known to occur when membrane-bound, the potential function of soluble LRP2 is unknown. In this work, we uncover a role for extracellular and soluble versus membrane-bound Lrp2 where proteolysis acts as a possible molecular switch for BMP activity. Functional roles depend on the protein state: full length Lrp2 facilitates intracellular signaling required to prevent ocular overgrowth, while the cleaved, extracellular domain antagonizes BMP signaling by acting as a ligand trap and promotes axial lengthening.

Myopia is the most prevalent visual disorder in the world (20,21), and one of the leading causes of blindness in many countries, especially in south-east Asia. High myopia (generally regarded as requiring more than −6.0 diopters of lens correction) elevates the risk of retinal detachment, cataract and glaucoma, as well as myopic macular degeneration, chorioretinal atrophy and staphyloma (22,23). At least 33% of adults in the US are affected by myopia (24), and preventing progression of refractive error to high myopia helps to reduce the occurrence of these secondary pathologies. There are a large number of genetic factors that contribute to onset of myopia, with many proteins associated with extracellular matrix (ECM), the visual cycle, and neuronal development (25,26). In addition, myopia is linked to environmental factors, where excessive time spent indoors without exposure to bright outdoor light levels appears to increase the risk of developing myopia (27,28). However, much remains unknown about the nature and components of signaling pathways that are dysregulated in myopia, as well as the cellular sites of action.

Unlike many anatomical structures, the eye has an absolute requirement for precise size control, since even small deviations in axial length have dramatic effects on refractive state. Small changes in axial length associated with genetic mutation or dysregulation of gene expression can have significant effects on refractive state, while only subtly affecting eye size. Continuous control of eye size to establish and maintain emmetropia after juvenile development has been demonstrated as necessary in all species studied (reviewed in (29)). Early post-natal eyes are ametropic, and visual cues inform the eye how to correct refractive errors. Signaling within the eye that mediates emmetropization is poorly understood, both in terms of the cells involved and the precise pathways utilized. Dysregulation of these pathways however, can lead to significant refractive error.

Mutations in BMP4 have been associated with a spectrum of ocular disorders, including high myopia. Individuals from families that frequently presented with anophthalmia-microphthalmia, retinal dystrophy, myopia, brain malformation and poly/syndactyly were found to have a nonsense frameshift mutation in BMP4 that segregated with these symptoms (30,31). Genome wide association studies (GWAS) identified linked DNA variants located near human *BMP2* and *BMP3* in myopic individuals (25,26). In animal studies, myopia induced by form-deprivation resulted in reduced BMP signals. In chicks, induced myopia led to a reduction in BMP2 in the retina/RPE (32), while in guinea pigs, form-deprivation myopia resulted in a reduction in BMP2 and BMP5 within the sclera (33,34). Similarly, defocus-induced myopia in guinea pigs caused a decrease in BMP2 in the sclera (35), and a decrease in BMP2/4/7 in the RPE (36,37). Conversely, lens-induced hyperopia has been shown to cause an increase in BMP2/4/7 in the chick RPE (36,37). Finally, prematurely-truncating mutations in the BMP antagonist Noggin causing NOG-related-symphalangism spectrum disorder/stapes ankylosis have been associated with hyperopia (38–40). Taken together, these data indicate a strong connection between BMP signaling and refractive error, where reduced BMP levels associate with myopia or lengthening of the eye, and increased BMP levels associate with hyperopia or shortening of the eye.

BMP ligands are members of the TGFβ superfamily whose receptors transduce signaling events from the cell surface to activate Smad and non-Smad signaling, as well as non-transcriptional responses (for review, see (41)). Prior to binding to the transmembrane receptors, BMPs can be bound by secreted, soluble ligand trap antagonists such as Noggin, which bind BMPs and prevent interaction with their receptors by acting as antagonists (42).

We have previously described a zebrafish *lrp2* mutant model, which shows enlarged eyes and high myopia, as well as elevated IOP and retinal ganglion cell damage (43,44). In this work, we profile gene expression in ocular tissues of wild-type and *lrp2*^-/-^ mutants at the onset of buphthalmia. We show that eye enlargement and myopia can be ameliorated in *lrp2*^-/-^ zebrafish mutants by expressing full-length Lrp2 in the RPE, indicating that Lrp2 acts locally within the eye to control emmetropization. We also demonstrate that Bmp4 binds to the first of the four LDL type A domains of Lrp2. To begin to address the role of cleaved and released extracellular Lrp2, we directed RPE-specific overexpression of Noggin3 (a high affinity BMP antagonist) and Gremlin2 (a lower affinity BMP antagonist). These studies showed that inhibiting BMP signaling leads to eye enlargement and myopia. Of particular relevance, we find that overexpressing a secreted version of the first LDL type A domain of Lrp2 from the RPE phenocopies the effects observed with the known BMP antagonists and exacerbates myopia in *lrp2* mutant zebrafish. We also demonstrate that extracellular cleavage of zebrafish Lrp2 occurs in cultured human cells, and find that cleavage is regulated by a beta-site amyloid precursor protein cleaving enzyme (BACE) protease. Furthermore, inhibition of BACE enzyme activity in the zebrafish eye increases Bmp signaling in the choroid/sclera, the primary region of extracellular remodeling associated with emmetropic growth. Similarly, expression of a constitutively soluble LDLA1 domain from the RPE reduces BMP activity, while expression of a membrane-bound form does not. Analysis of *lrp2*^-/-^ and wild-type eye transcriptomes shows altered expression of ECM components. We propose that modulation of Bmp signaling, facilitated by Lrp2 and its cleavage state, contributes to controlling eye size and prevents aberrant growth and myopia.

## Results

### Loss of Lrp2 results in alterations to multiple signaling pathways within the eyes

Loss of function mutations of *lrp2* lead to enlarged, myopic eyes in zebrafish and other species (6,7,43,45). However, the effect of disrupting Lrp2 activity on specific signaling pathways and cellular processes has not been extensively assessed in ocular tissues. Because ocular histology and growth is normal at 1 mpf and earlier in *bugeye* fish (46), we performed transcriptome analyses at this stage to avoid potential confounding secondary and/or pathology-related pathway changes that occur in more aged animals. Specifically, we isolated RNA from 1 mpf wild-type and *lrp2*^-/-^ eyes, as well as dissected ocular tissues (sclera/choroid, RPE, retina), where each condition was replicated in triplicate with independent biological samples. RNA extracted from these samples was assayed for tissue specificity using retinal-, RPE- or sclera/choroid-specific marker genes by qPCR (S1 Figure), and then processed for deep RNA sequencing (RNAseq) (S1 Table, S2 Table). Subsequent bioinformatic analysis was applied to detect potential changes in signaling pathways associated with loss of Lrp2.

Using curated databases, such as Gene Ontology, DAVID, KEGG, and IPA, for transcript signatures associated with different types of signaling, we assessed whether genes altered by loss of Lrp2 correlated with those of specific pathways. Analyses indicated that Bmp and Shh signaling were both altered in *lrp2* mutant eyes (S3 Table). However, expression signatures associated with these databases are assembled from published data derived from non-ocular tissues of non-zebrafish origin. Since Bmp and Shh signaling pathways are strongly associated with control of eye size and development, and both have been previously shown to be regulated by Lrp2, we generated ocular-specific, zebrafish transcript signatures for Bmp and Shh. This was accomplished by using transgenic zebrafish lines to acutely overexpress either Shha or Bmp4 under inducible heat-shock control. At 5 dpf, lines *hsp70l:shha*-*eGFP* (47), or *hsp70l:eGFP*-*bmp4* (this study) were heat-shocked before isolation of GFP-positive eyes 8 hours later. RNA expression profiles from these eyes were used to define zebrafish ocular responses to Shh and Bmp pathway activation (S2 Figure, S3 Figure).

These signatures were used to carry out gene-set enrichment analysis (GSEA) and assess overlap of the gene sets in Lrp2-deficient ocular tissues (see below). GSEA examines transcriptome datasets to determine whether a defined list of genes (referred to as signatures or commonly-regulated gene sets) is concordantly and statistically altered between control and experimental groups (48,49). Specifically, RNA profiles of whole eye and dissected ocular tissues from 1 mpf wild-type and *lrp2*^-/-^ mutant zebrafish were compared to eye-specific Bmp4 and Shh reference gene sets. Both pathways were significantly altered in the absence of Lrp2. To prioritize the gene pathway and eye tissue of greatest importance, we applied a simple scoring system to assess the significance of Lrp2 for each pathway in specific eye tissues (Table 1). For each sample assessed, nominal GSEA p-values lower than 0.05 were scored as +1, while p-values greater than 0.05 were scored as 0. The final scores were summed and used to rank significance of each signaling pathway in each tissue. We found the Shh pathway was broadly affected and scored highest for whole eyes. Bmp pathway genes were more selectively altered within the RPE, but also showed changes in the sclera/choroid and neural retina. Because Shh has already been studied in the context of eye size regulation using mice (9), and RPE is known as a key signaling tissue for emmetropization and thought to mediate signals processed in the sclera and choroid, we concentrated our analysis on Bmp signaling associated with those tissues.

**Table 1.** Gene set enrichment analysis of wild-type and *lrp2*^-/-^ zebrafish eyes and dissected tissues using Bmp4 and Shha pathway signature sets from this work. GSEA analyses were conducted on RNAseq datasets from whole eye and dissected eye tissues using custom BMP4 and Shha-specific signatures, p-values lower than 0.05 were scored as +1, while p-values greater than 0.05 were scored as 0. The final scores were summed and used to rank significance of each signaling pathway in each tissue.

| Tissue | Pathway | Bmp4 | Bmp4 | Shha | Shha | Nominal score |  |
| --- | --- | --- | --- | --- | --- | --- | --- |
| Whole eye | Direction of regulation | Up | Down | Up | Down | Bmp4 | Shha |
| Upregulated in class |  | *lrp2* | wild-type | wild-type | *lrp2* |  |  |
| Enrichment Score (ES) |  | 0.272 | -0.343 | -0.469 | 0.353 |  |  |
| Normalized Enrichment Score (NES) |  | 1.183 | -1.410 | -1.887 | 1.415 |  |  |
| Nominal p-value |  | 0.114 | 0.024 | 0.000 | 0.020 |  |  |
| FDR q-value |  | 0.114 | 0.024 | 0.000 | 0.020 |  |  |
| FWER p-Value |  | 0.087 | 0.008 | 0.000 | 0.014 |  |  |
| Score |  | 0 | +1 | +1 | +1 | +1 | +2 |
| Tissue | Pathway | Bmp4 | Bmp4 | Shha | Shha |  |  |
| Sclera/choroid | Direction of regulation | Up | Down | Up | Down |  |  |
| Upregulated in class |  | *lrp2* | *lrp2* | wild-type | *lrp2* |  |  |
| Enrichment Score (ES) |  | 0.263 | 0.457 | -0.383 | 0.381 |  |  |
| Normalized Enrichment Score (NES) |  | 1.121 | 1.712 | -1.680 | 1.489 |  |  |
| Nominal p-value |  | 0.221 | 0.002 | 0.010 | 0.011 |  |  |
| FDR q-value |  | 0.221 | 0.002 | 0.010 | 0.011 |  |  |
| FWER p-Value |  | 0.205 | 0.002 | 0.002 | 0.009 |  |  |
| Score |  | 0 | +1 | +1 | +1 | +1 | +2 |
| Tissue | Pathway | Bmp4 | Bmp4 | Shha | Shha |  |  |
| RPE | Direction of regulation | Up | Down | Up | Down |  |  |
| Upregulated in class |  | *lrp2* | wild-type | wild-type | *lrp2* |  |  |
| Enrichment Score (ES) |  | 0.353 | -0.327 | -0.507 | 0.337 |  |  |
| Normalized Enrichment Score (NES) |  | 1.598 | -1.408 | -2.130 | 1.387 |  |  |
| Nominal p-value |  | 0.001 | 0.017 | 0.000 | 0.032 |  |  |
| FDR q-value |  | 0.001 | 0.017 | 0.000 | 0.032 |  |  |
| FWER p-Value |  | 0.001 | 0.005 | 0.000 | 0.024 |  |  |
| Score |  | +1 | +1 | +1 | +1 | +2 | +2 |
| Tissue | Pathway | Bmp4 | Bmp4 | Shha | Shha |  |  |
| retina | Direction of regulation | Up | Down | Up | Down |  |  |
| Upregulated in class |  | wild-type | wild-type | wild-type | *lrp2* |  |  |
| Enrichment Score (ES) |  | -0.256 | -0.241 | -0.419 | 0.338 |  |  |
| Normalized Enrichment Score (NES) |  | -1.181 | -0.969 | -1.607 | 1.425 |  |  |
| Nominal p-value |  | 0.074 | 0.499 | 0.012 | 0.017 |  |  |
| FDR q-value |  | 0.074 | 0.499 | 0.012 | 0.017 |  |  |
| FWER p-Value |  | 0.041 | 0.270 | 0.006 | 0.008 |  |  |
| Score |  | +1 | 0 | +1 | +1 | +1 | +2 |

### RPE-driven expression of Lrp2 partially rescues the buphthalmic *bugeye* phenotype

To investigate whether buphthalmia was caused by absence of Lrp2 from the eyes, which is expressed in the RPE, or by systemic changes in circulating ligands normally bound by Lrp2 in the kidney and elsewhere, we developed an RPE-specific promoter to express Lrp2 only in that tissue. Similar to studies using mammalian *RPE65*, we isolated approximately 800 bp upstream of the translational start site of zebrafish *rpe65a* (50). We found that this promoter drove fluorescent transgene expression in developing and adult RPE and pineal (Figure 1). In embryos, *rpe65a:eGFP* transgenic expression was broad, but low. In larvae, we observed strong enrichment of fluorescent expression in developing RPE cells starting just prior to when they resolve to a monolayer (51). Subsets of cells within the pineal also showed enriched eGFP expression. In adult zebrafish, eGFP expression was maintained and restricted to the RPE and pineal.

**Figure 1.**
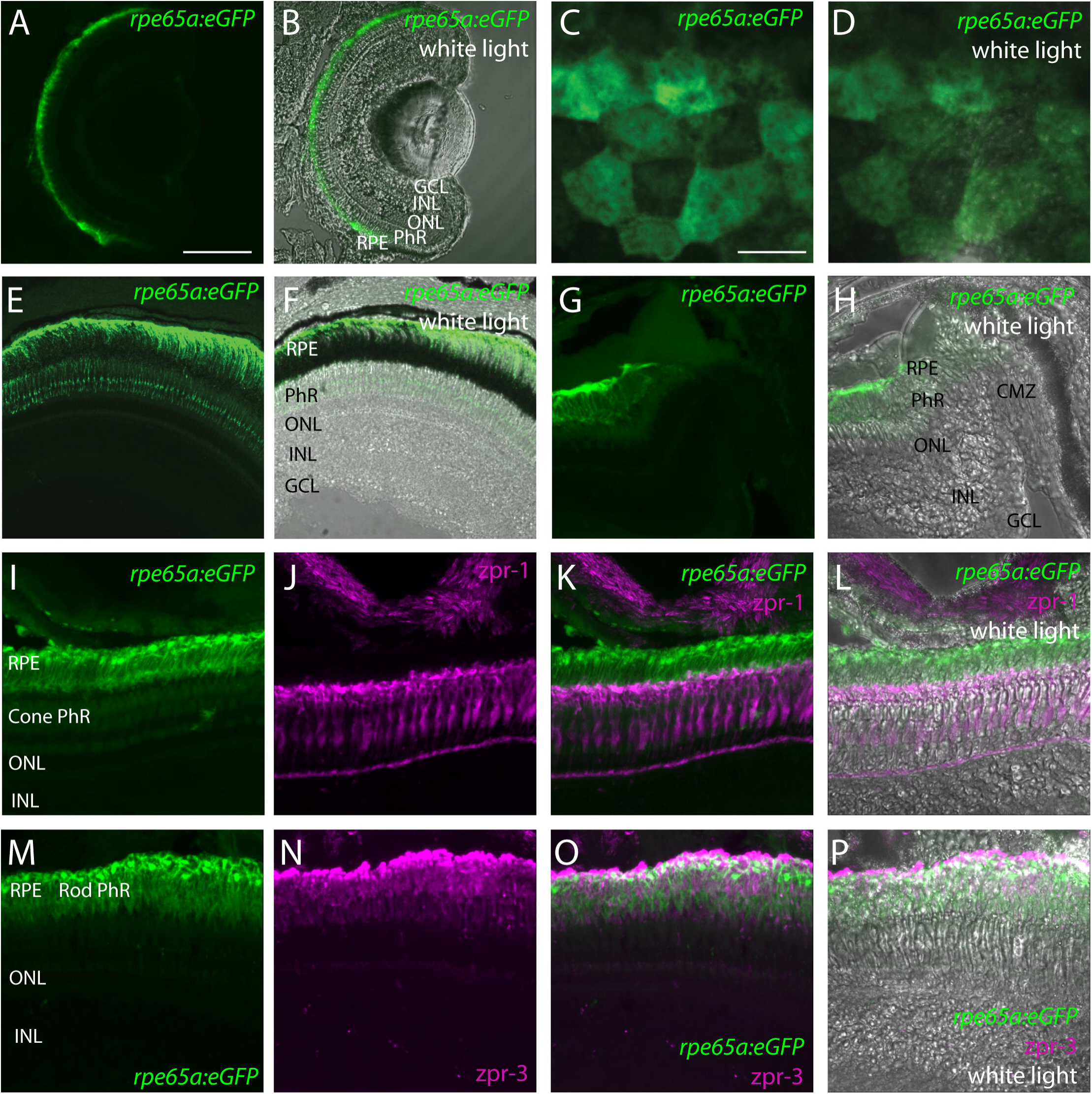
The zebrafish *rpe65a* promoter drives transgene expression in the developing and adult RPE. A - D. the *rpe65a* promoter drives expression in the 5 dpf RPE (A,B, cryosection; C, D, en face in vivo). E - H. Strong eGFP expression is seen in the 3 mpf adult retina but not the CMZ. I -L. RPE expression (green) driven by the *rpe65a* promoter contrasts with red-green cone-specific zpr-1 antibody staining (magenta). M – P. Similarly, eGFP in RPE cells contrast with rod-specific zpr-3 antibody staining (magenta). Scale bars: A, B, E – P = 50 μm; C, D = 10 μM. RPE, retinal pigment epithelium; PhR, photoreceptor; ONL, outer nuclear layer; INL, inner nuclear layer; GCL, ganglion cell layer; CMZ, ciliary marginal zone.

We next isolated the full-length zebrafish *lrp2* coding sequence and generated transgenic lines expressing both *rpe65a:Gal4* and DsRed ←*UAS* →Lrp2-HA (abbreviated to *rpe65a>lrp2*-*HA*). This transgenic line uses the *rpe65a* promoter to drive Gal4 expression, which transactivates the upstream activating sequence (UAS) bi-directionally so that fluorescent DsRed marks cells co-expressing Lrp2-FIA. At 3 mpf, wild-type and *lrp2*^-/-^ fish with or without RPE-driven Lrp2-FIA expression were examined using spectral domain - ocular coherence tomography (SD-OCT) to assess eye size relative to body axis and calculate the relative refractive error, a measure of deviation from emmetropia. This imaging technique allows for precise measurements in live samples, avoiding tissue fixation artifacts and enabling detection of the small differences in axial length that have major impact on visual blur (52). Adult *lrp2*^-/-^ mutant zebrafish carrying the *rpe65a>lrp2*-*HA* transgenes had significantly smaller eyes (lower eye axial length: body axis ratios) than non-transgenic, mutant siblings (Figure 2 A). Flowever, *lrp2*^-/-^ *rpe65a>lrp2*-*HA* eyes still had significantly larger eyes than wild-type controls. Likewise, *lrp2*^-/-^ adult zebrafish carrying the *rpe65a>lrp2* transgenes had significantly reduced relative refractive errors (less myopic) when compared to non-transgenic, mutant siblings (Figure 2 B). Similar changes were measured in replicate experiments. Furthermore, expressing Lrp2 directly from the *rpe65a* promoter also resulted in abatement of enlarged eyes and myopia (data not shown). Importantly, the *rpe65a* promoter does not show activity within the ciliary margin zone (Figure 1 G), and so the rescue measured is due only to RPE cell expression and not from the ciliary region, which is known to have strong Shh activity. In summary, this data indicate that RPE-expressed Lrp2 contributes to regulation of eye growth and emmetropization.

**Figure 2.**
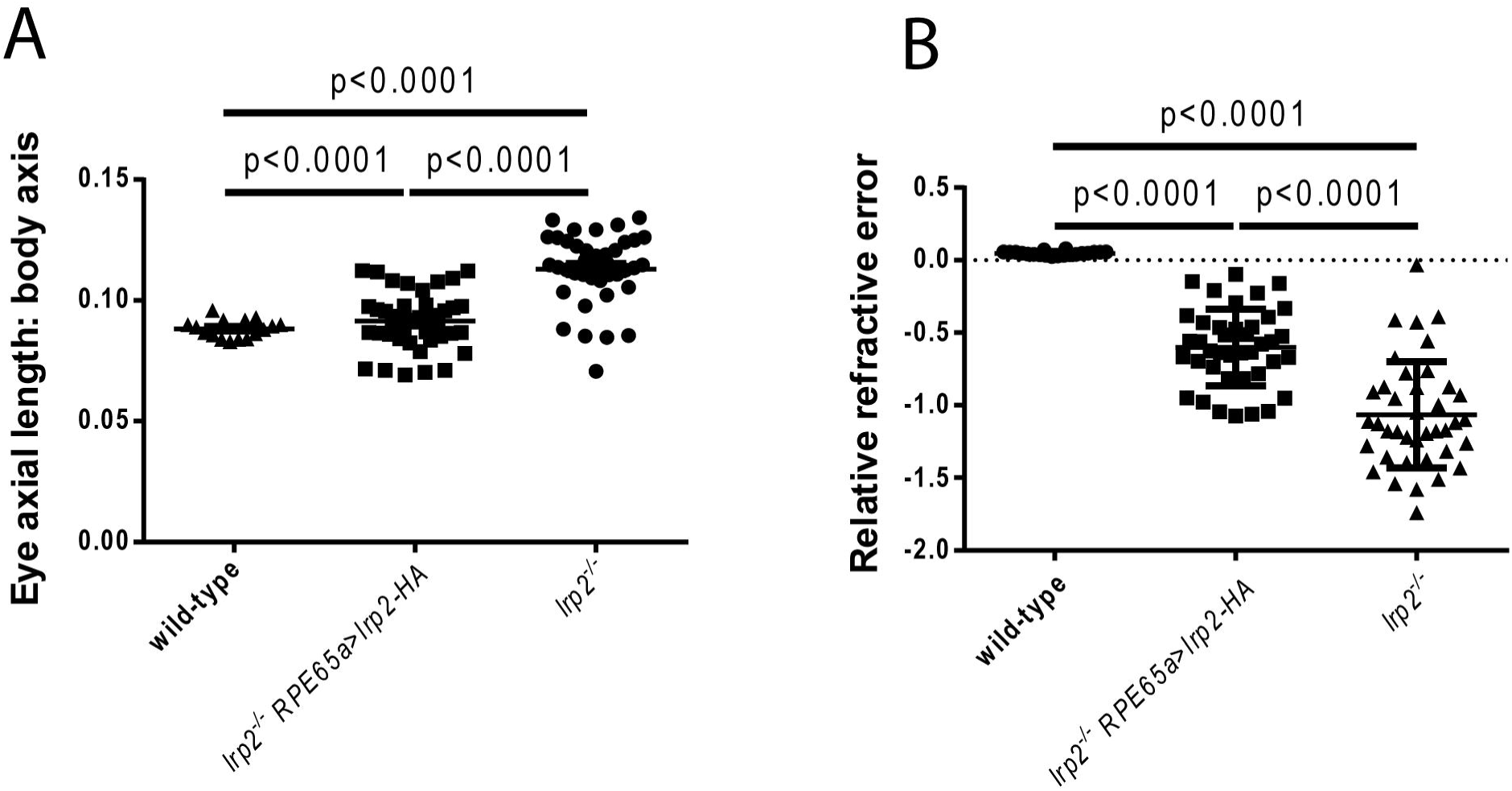
*rpe65a*-driven Lrp2 partially rescues the myopic *lrp2/bugeye* phenotype in adult zebrafish. A. Eye axial length: lens diameter ratio is reduced in *lrp2*^-/-^ expressing RPE-derived Lrp2 at 3 mpf (0.09143 vs 0.1130; p=0.0001; Kruskal-Wallis test). Flowever, *lrp2*^-/-^ *rpe65a>lrp2*-*HA* eyes still have a significantly greater eye axial length: lens diameter ratio than wild-type controls (0.088 vs 0.1130; p<0.0001; Kruskal-Wallis test). B. Similarly, the relative refractive error is lessened in *lrp2*^-/-^ eyes expressing RPE-derived Lrp2 at 3 mpf (−0.6013 vs −1.066; p<0.0001; Kruskal-Wallis test). This rescue is partial and does not restore myopic *lrp2*^-/-^ eyes to emmetropia (0.048 vs −1.066; p<0.0001; Kruskal-Wallis test). This experiment was performed twice for validation with similar results.

### eGFP-Bmp4 is functional and perdures in discrete locations during development

Because eye size and refraction were rescued by RPE-expression of Lrp2 within *lrp2* mutants, and *lrp2* mutant RPE showed defects in BMP signaling, we sought to explore more directly the relationship of Lrp2 and BMP activity using an eGFP-tagged Bmp4. To validate that eGFP-Bmp4 was functional, we generated a transgenic line, *hsp70l:eGFP*-*bmp4*, that placed the Bmp4 fusion protein downstream of a heat shock-inducible promoter, allowing activation of eGFP-Bmp4 expression at various times during development. Fleat-induced eGFP-Bmp4 expression was verified by Western blot (S4 Figure). Secretion and activity was confirmed by inducing global eGFP-Bmp4 expression at 30 – 50% epiboly, which resulted in extracellular eGFP and severe ventralization phenotypes, similar to the patterning defects observed in zebrafish following overexpression of native Bmp at these early times (53).

At embryonic stages after axis patterning, expression of eGFP-Bmp4 resulted in milder phenotypes. Expression induced at 1, 2, 3 or 4 dpf followed by image analysis 24 hours later revealed patterns of perdurance of eGFP-Bmp4. Stabilized eGFP-Bmp4 was enriched in cells of the eye, head, somites and other areas (S4 Figure). Despite the mild phenotypic consequences at later stages, eGFP-Bmp4 expression did stimulate Smad signaling, as shown by elevated red fluorescence from the BRE:dmKO2 reporter (54). These data verify the bioactivity of eGFP-Bmp4.

Given the association between Bmp4 levels and refractive state of the eye, we attempted to overexpress Bmp4 from the RPE. To do so, we generated a *UAS:eGFP*-*bmp4* transgenic line to combine with the *rpe65a:Gal4* transgene and achieve sustained RPE-derived overexpression of eGFP-Bmp4. However, *rpe65a* and our promoter transgene expresses broadly during early development before it becomes confined to the RPE and pineal, resulting in severe microphthalmia ((55), this study). Interestingly, these embryos often developed into adults with abnormal phenotypes restricted to the eyes, including coloboma and disorganization of the retina (S5 Figure). Since the eyes did not develop correctly, we were unable to use this strategy to isolate the effects of Bmp4 overexpression on ocular size and the refractive state.

### Overexpression of BMP ligand-trap antagonists from the RPE alter eye size and cause myopia

To test whether BMP activity associated with the RPE is critical for eye growth and emmetropization, we overexpressed the potent Bmp antagonist zebrafish Noggin3 (Nog3) from that epithelium. We initially tried to create a double transgenic line that bi-directionally expresses *DsRed*← *UAS*→*nog3* within a stable *rpe65a:Gal4* line. Despite repeated attempts, we were unable to establish founder fish that gave rise to offspring transgenic for both constructs, likely due to deleterious effects of expressing Noggin3 broadly during early development, as was the case for Bmp4 overexpression (S5 Figure). Instead, we injected the plasmid carrying the *DsRed*←*UAS*→*nog3* sequence into *rpe65a:Gal4* embryos (either *lrp2*^+/-^ as sibling controls, or *lrp2*^-/-^ mutants) and screened for red fluorescence expression in RPE cells at 5 dpf. These fish, abbreviated *rpe65a>nog3*, were raised to adulthood, and eye and body metrics measured at 2 months.

Overexpression of Nog3 in RPE of *lrp2*^-/-^ zebrafish exacerbated the enlarged eye phenotype and enhanced the degree of myopia as compared to non-transgenic siblings that were also mutant for *lrp2* (Figure 3 A, B). Interestingly, overexpression of Nog3 in *lrp2*^+/-^ zebrafish, which are indistinguishable from wild-type fish, induced eye enlargement and refractive errors. These results show that RPE-based overexpression of Nog3 increases the degree of myopia in *lrp2*^-/-^ adult zebrafish, and causes myopia in *lrp2*^+/-^ zebrafish that are otherwise slightly hyperopic.

**Figure 3.**
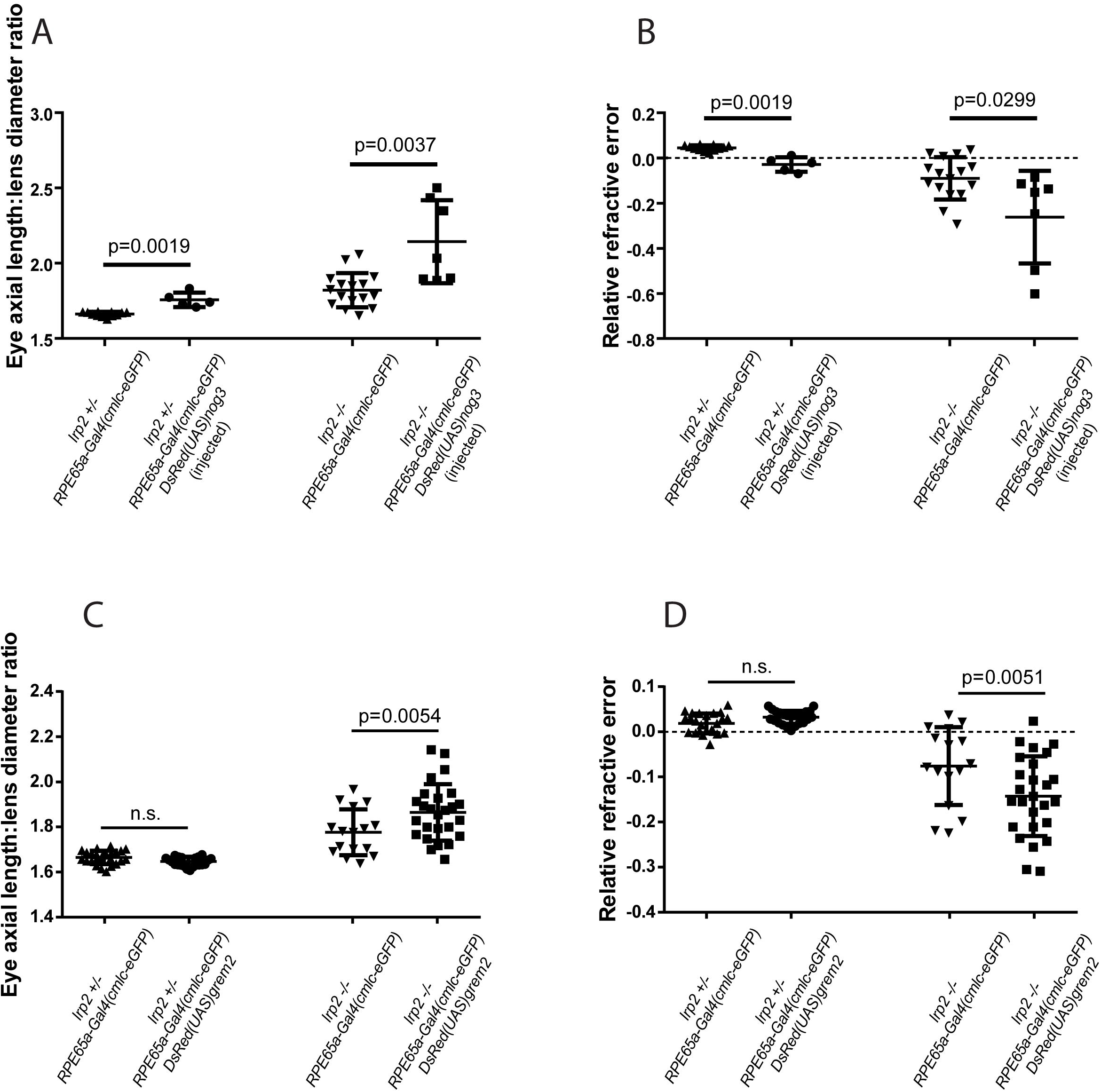
*rpe65a*-driven Noggin3 expression causes myopia in wild-type zebrafish and exacerbates myopia in *lrp2/bugeye* adults, while *rpe65a*-driven Gremlin2 expression exacerbates myopia in *lrp2*^-/-^ but not wild-type. A. RPE-derived Nog3 causes an increase in axial length: lens diameter ratio in both *lrp2*^-/-^ and wild-type zebrafish at 2 mpf (*lrp2*^-/-^ 2.14 vs 1.82; p=0.0037; Mann-Whitney test. *lrp2*^-/-^: 1.76 vs 1.66; p=0.0019; Mann-Whitney test. n= 5 for *lrp2*^+/^~ *rpe65a>nog;* n= 7 for *lrp2*^-/-^ *rpe65a*>*nog;* n = 12 for *lrp2*^-/-^ *rpe65a:Gal4*; n = 16 for *lrp2*^-/-^ *rpe65a:Gal4*). B. RPE-derived Nog3 expression worsens the degree of relative refractive error in both *lrp2*^-/-^ and wild-type zebrafish at 2 mpf (*lrp2*^-/-^: 0.262 vs 0.049; p=0.0299; Mann-Whitney test. *lrp2*^+/-^: −0.029 vs 0.044; p=0.0019; Mann-Whitney test.). C. RPE-derived Grem2 causes an increase in axial length: lens diameter ratio in *lrp2*^-/-^ zebrafish at 2 mpf, but not in wild-type (*lrp2*^-/-^: 1.87 vs 1.78; p=0.0054; Mann-Whitney test, n was greater than 24 for each genotype, except for *lrp2*^-/-^ *rpe65a:Gal4*, where n = 16.). D. RPE-derived Grem2 expression worsens the degree of relative refractive error in *lrp2*^-/-^ zebrafish at 2 mpf, but not wild-type (*lrp2*^-/-^: −0.143 vs −0.076; p=0.0051; Mann-Whitney test.).

To complement the Noggin3 studies and attempt to generate stable transgenic fish capable of expressing a BMP antagonist, we tested the effects of a weaker BMP-binding inhibitor, Gremlin2, on eye size and refractive state. Indeed, we were able to generate double transgenic lines of *rpe65a:Gal4; DsRed*←*UAS*→*grem2* (abbreviated to *rpe65a>grem2*). These fish overexpress Grem2 from the RPE, and show normal survival and fertility. Overexpression of Grem2 from *lrp2*^-/-^ RPE cells increased the relative ocular axial length (Figure 3 C). The mean relative refractive error was also significantly affected (Figure 3 D). Flowever, in *lrp2*^+/-^ zebrafish, overexpression of Grem2 did not significantly alter eye size or impact the refractive state. Cumulatively, these data are consistent with the concept that modulation of BMP signaling within the RPE and adjacent tissues can regulate ocular growth and emmetropization.

### Bmp4 binds to the LDLA1 domain of Lrp2 in vivo

Bmp4 has previously been shown to bind purified Lrp2 in vitro, though the site of interaction has not been discovered, and direct interaction has not been verified on cells (56). The extracellular portion of Lrp2 can be divided into four domains, characterized by LDLA motifs. We generated four expression vectors corresponding to the isolated LDLA domains 1-4, and individually fused each domain with an S•tag epitope in plasmids CMV/SP6:LDLAn-S•tag, where n corresponds to LDLA domain 1, 2, 3 or 4. In addition we constructed an epitope-tagged Bmp4 to investigate whether, and to which domain of Lrp2, BMP4 binds in vivo (S4 Figure).

Zebrafish Lrp2 LDLA domains 1 through 4 were transfected along with eGFP-tagged Bmp4 into HEK293T cells. LDLA domains were immobilized from conditioned medium and extracted cellular proteins before probing for bound eGFP-Bmp4 by Western blot. Comparing total input protein against protein retained after immunoprecipitation showed that eGFP-Bmp4 binds to the LDLA1 domain, but not to LDLA domains 2, 3 or 4 (Figure 4). Importantly, binding of ligands to extracellular regions of Lrp2, which may be cleaved from the cell surface, suggests that Lrp2 might act as a ligand trap as well as an endocytic receptor.

**Figure 4.**
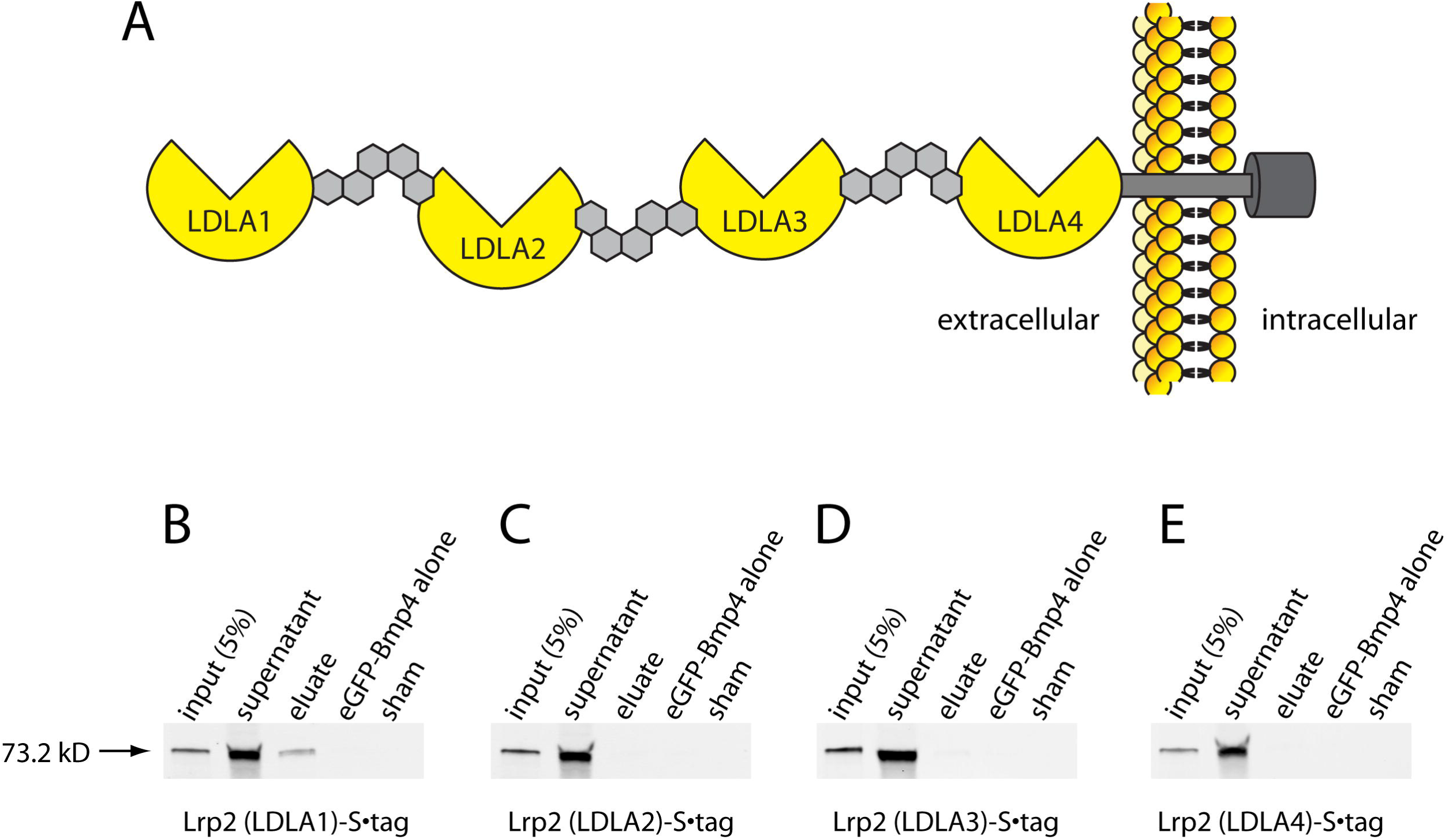
The first LDLA domain of Lrp2 binds Bmp4 *in vivo*. A. Schematic showing the structure of Lrp2 with four LDLA domains that bind cognate ligands. B – E. Expression of individual LDLA domains 1 – 4 with eGFP-Bmp4 shows that immobilized LDLA1-S•tag can bind Bmp4, but that the other LDLA domains cannot.

### RNAseq shows altered gene signaling in dissected eye tissues of *lrp2*^-/-^ and wild-type

To evaluate which Bmp ligands and receptors are available for signaling in the RPE and adjacent tissues, we evaluated RNAseq data from wild-type 1 mpf eyes and dissected tissues (S6 Figure). Most Bmp ligands were more highly expressed in RPE and choroid/sclera than retina or whole eye (one-way ANOVA with Tukey’s multiple comparison test), though we note that *bmpr2a*, *bmpr2b* and *acvr2aa* are more highly expressed in the retina than elsewhere. The Bmp receptors *bmpr1ab* and *bmpr1ba* were more highly expressed in the RPE than other tissues, and *bmpr1ab* was also enriched in the sclera/choroid. Activin receptors, which can also transduce Bmp signals as well as other TGFβ superfamily signals, were more broadly expressed in all eye tissues, though *acvrl1/alk1* and *acvrl1/alk2/alk8* were enriched in the RPE. We note that Bmp type II receptors (*bmpr2a*, *bmpr2b*) can complex with Alk1 (AcvrL1) and Alk2 (Acvr1) as well as Bmp type I receptors, and it may be that TGFβ signaling unrelated to Bmp signaling is the reason for high levels of *bmpr2a* and *bmpr2b* in the retina. Taken together, these data suggest that Bmps are important for signaling between the RPE and sclera/choroid, consistent with a model in which Bmp activity mediates the ability of RPE to direct emmetropic remodeling of the sclera/choroid.

### Secretion of the LDLA1 domain of Lrp2 from the RPE increases myopia in an *lrp2*^-/-^ zebrafish

Because full-length Lrp2 overexpression reduced the large eye phenotype in *lrp2*^-/-^ zebrafish, and Bmp4 bound only to LDLA1, we wanted to test whether expression of the LDLA1 domain alone from the RPE was sufficient to rescue the *bugeye* phenotype. We placed DNA encoding the LDLA1 domain downstream of the native Lrp2 signal peptide to ensure proper export of the resulting protein. Using this plasmid, we made the following transgenic line: *DsRed*←*UAS*→*lrp2,signal peptide*-*LDLA1* (abbreviated to *rpe65a>LDLA1*). These fish were combined with *rpe65a:Gal4* carriers to overexpress the secreted N-terminal domain of Lrp2 (signal peptide-LDLA1) from the RPE. Double transgenic progeny grew to maturity and were fertile. Overexpression of the Lrp2 LDLA1 domain from the RPE of *lrp2*^-/-^ mutants increased both the normalized eye size and relative refractive error compared to mutant siblings that only expressed Gal4 (Figure 5 A, B). However, overexpression of LDLA1 did not significantly affect the eyes of *lrp2*^+/-^ zebrafish. These results show that overexpression of RPE-secreted Lrp2 LDLA1 domain increases the degree of myopia in *lrp2*^-/-^ adult zebrafish, similar to the results found with overexpression of either Nog3 or Grem2.

**Figure 5.**
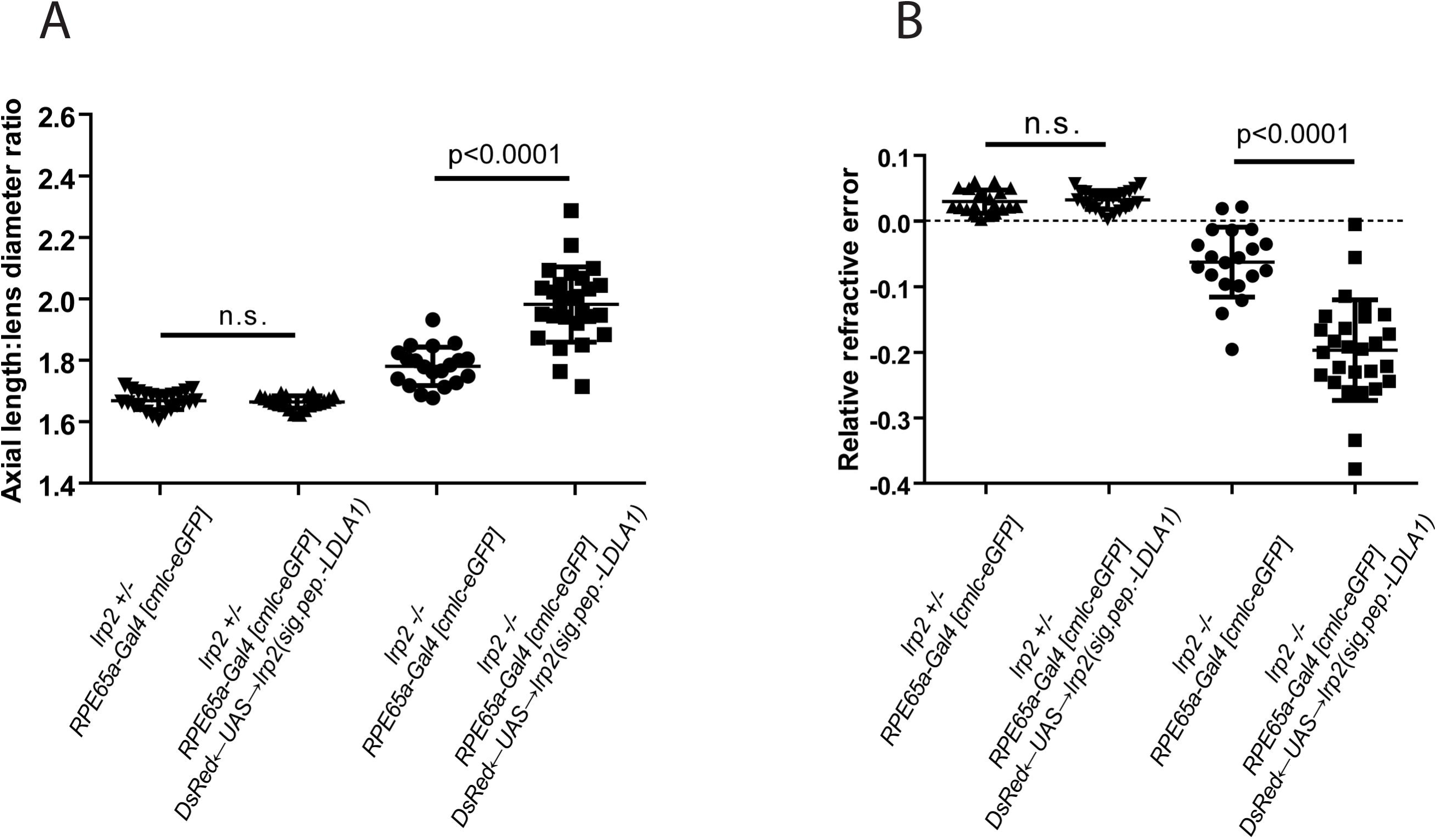
*rpe65a*-driven soluble Lrp2 (LDLA1) exacerbates myopia in *lrp2/bugeye* adults. A. RPE-derived Lrp2 (LDLA1) causes an increase in axial length: lens diameter ratio in *lrp2*^-/-^ zebrafish at 2 mpf, but not in wild-type (*lrp2*^-/-^: 1.98 vs 1.78; p<0.0001; Mann-Whitney test, n = greater than 20 for each genotype.). B. RPE-derived Lrp2 (LDLA1) expression worsens the degree of relative refractive error in *lrp2*^-/-^ zebrafish at 2 mpf, but not wild-type(*lrp2*^-/-^: −0.197 vs −0.063; p<0.0001; Mann-Whitney test.).

### Lrp2 is cleaved extracellularly to release a soluble domain

Replacement of full-length Lrp2 in the *lrp2* mutant RPE reduces the severity of the buphthalmic phenotype; however, overexpression of the secreted, Bmp4-binding domain exacerbates eye enlargement. This suggests that soluble Lrp2 has a different function versus membrane-bound Lrp2, and that release of the extracellular domain by proteolytic cleavage may act as a functional switch. In the eye, proteolysis of Lrp2 may serve to fine-tune signaling that controls emmetropization. In cultured proximal tubule cells of the kidney, LRP2 has been shown to be cleaved to produce a soluble extracellular N-terminal domain and an intracellular C-terminal domain. To examine the possibility of extracellular cleavage of zebrafish Lrp2, and investigate the factors that modulate this event, we generated a plasmid that placed secreted eGFP (seGFP) upstream of a 691 amino acid fragment of Lrp2 (457 a.a. extracellular domain + 234 a.a. of the full transmembrane and intracellular domains), and placed mCherry at the C-terminus to make *seGFP*-*lp2 (ECD/ICD)*-*mCherry* (Figure 6 A). This dually-tagged *lrp2* fragment was put under the control of a CMV promoter and expressed in HEK293T cells. Conditioned culture medium was harvested after 48 hours following transfection of the plasmid and the Lrp2 epitopes were probed by Western blot. As controls, *seGFP*-*V2A*-*mCherry* (constitutive cleavage) and *seGFP*-*mCherry* (no cleavage) were also placed under the control of the CMV promoter and transfected into equivalent batches of cells.

**Figure 6.**
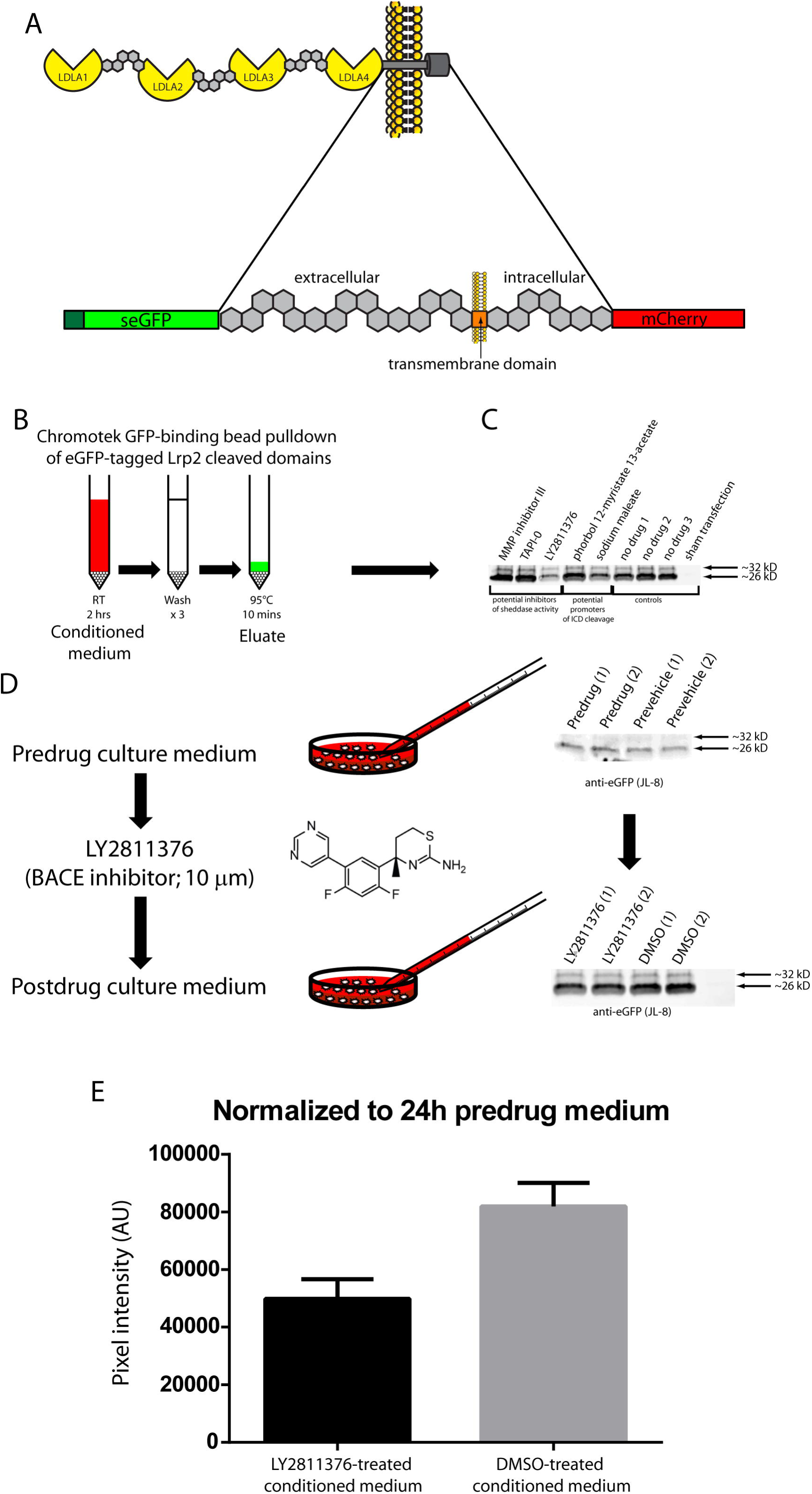
Lrp2 is cleaved extracellularly by a BACE family member to release a soluble form. A. Schematic showing tagging of a 432 amino acid region of Lrp2 with secreted eGFP at the N-terminus and mCherry at the C-terminus to monitor extracellular cleavage, referred to as seGFP-Lrp2 (ECD/ICD)-mCherry. B - D. HEK293T cells transfected with seGFP-Lrp2 (ECD/ICD)-mCherry were treated 24 hours after transfection with compounds and grown for another 24 hours before harvesting conditioned medium. eGFP-tagged domains cleaved from seGFP-Lrp2 (ECD/ICD)-mCherry were immobilized using Chromotek GFP-binding agarose before Western blotting and detection with JL-8 (anti-GFP antibody). E. When soluble eGFP-species levels from post-drug treatments were normalized to pre-treatment levels from the same culture plates, LY2811376-treatment appeared to reduce extracellular cleavage of seGFP-Lrp2 (ECD/ICD)-mCherry. Pixel intensity is expressed in arbitrary units based on ImageJ measurements of Western blot band densities.

To investigate regulation of extracellular cleavage, and to gain insight into the class of proteases that might mediate cleavage, we used pharmacological agents known to affect the activity of different classes of sheddases. The plasmid *CMV:seGFP*-*Lrp2 (ECD/ICD)*-*mCherry* was transfected into HEK293T cells for 24 hours, before cells were treated with a variety of compounds that either inhibit or promote extracellular cleavage of LDL receptor family members. To inhibit sheddases, matrix metalloprotease inhibitor III, TAPI-0 or LY2811376 were used to block activity of matrix metalloproteases, TNF-α converting enzyme (TACE)/ADAM17, and BACE1, respectively. To augment sheddase activity we added phorbol 12-myristate 13-acetate (PMA) and sodium maleate, which are known to stimulate cleavage of amyloid precursor protein. Transfected cells were treated with compounds for 24 hours before conditioned medium was collected. Conditioned medium from each group was applied to anti-GFP-conjugated agarose to bind any Lrp2 that may have been released into the medium through cleavage (seGFP-Lrp2 ECD), or potentially exocytosis or cell lysis (*seGFP*-*Lrp2 (ECD/ICD)*-*mCherry*) (Figure 6B). Bound proteins were eluted from the agarose and detected by Western blotting using anti-eGFP antibody (JL-8). Of the compounds tested, only LY2811376 altered the amount of eGFP-containing proteins in the medium, suggesting that BACE1 (or a similar enzyme that is also inhibited by LY2811376) mediates extracellular cleavage of Lrp2 (Figure 6 C). To further confirm BACE1-mediated cleavage of Lrp2, the experiment was repeated in duplicate, with culture medium collected before (24 hours after transfection) and after (48 hours after transfection) drug administration (Figure 6 D). Following LY2811376 treatment, when the amount of eGFP-containing proteins was normalized to pre-drug levels, there was an ~40% average reduction in extracellular cleavage observed (Figure 6 E). These results indicate that release of the extracellular region of zebrafish Lrp2 is mediated in part by a BACE1-like activity, and the responsible protease(s) may modulate Lrp2 function in vivo. Consensus substrate sites for BACE enzymes have been published, and comparisons with zebrafish Lrp2 protein sequence suggests several sites for BACE enzyme activity (S7 Figure). Our results so far suggest that BACE enzyme(s) are capable of extracellular cleavage of ectopically expressed Lrp2 and the released LDLA1 domain can bind Bmp and affect signaling.

### Inhibition of BACE activity increases Bmp signaling in tissues basal to the RPE

We next tested whether inhibiting BACE activity affects Bmp signaling in vivo using zebrafish. To inhibit BACE activity, we used two potent compounds, LY2811376 and BACE inhibitor II. Larval zebrafish transgenic for the Bmp-response element:d2GFP (*BRE:d2GFP*) express destabilized GFP to report the location and level of Bmp signaling in vivo (57). *BRE:d2GFP* larvae were treated at 4 dpf for 24 hours with a) 150 μm LY2811376 (BACE1 inhibitor), b) 50 μm BACE inhibitor II, or c) DMSO vehicle as control. At 4 dpf, zebrafish eyes show highest Bmp activity in the outermost layers (RPE, choroid, sclera) (Figure 7A). Following drug or vehicle treatment, eyes of *BRE:d2GFP* larvae were imaged by fluorescent microscopy and the signal in the outer layers of the eye was quantified. BACE inhibitor drug treatments led to higher levels of Bmp signaling in outer layers of reporter zebrafish when compared to vehicle controls for both LY2811376 and BACE inhibitor II (Figure 7 A, B, F, G).

**Figure 7.**
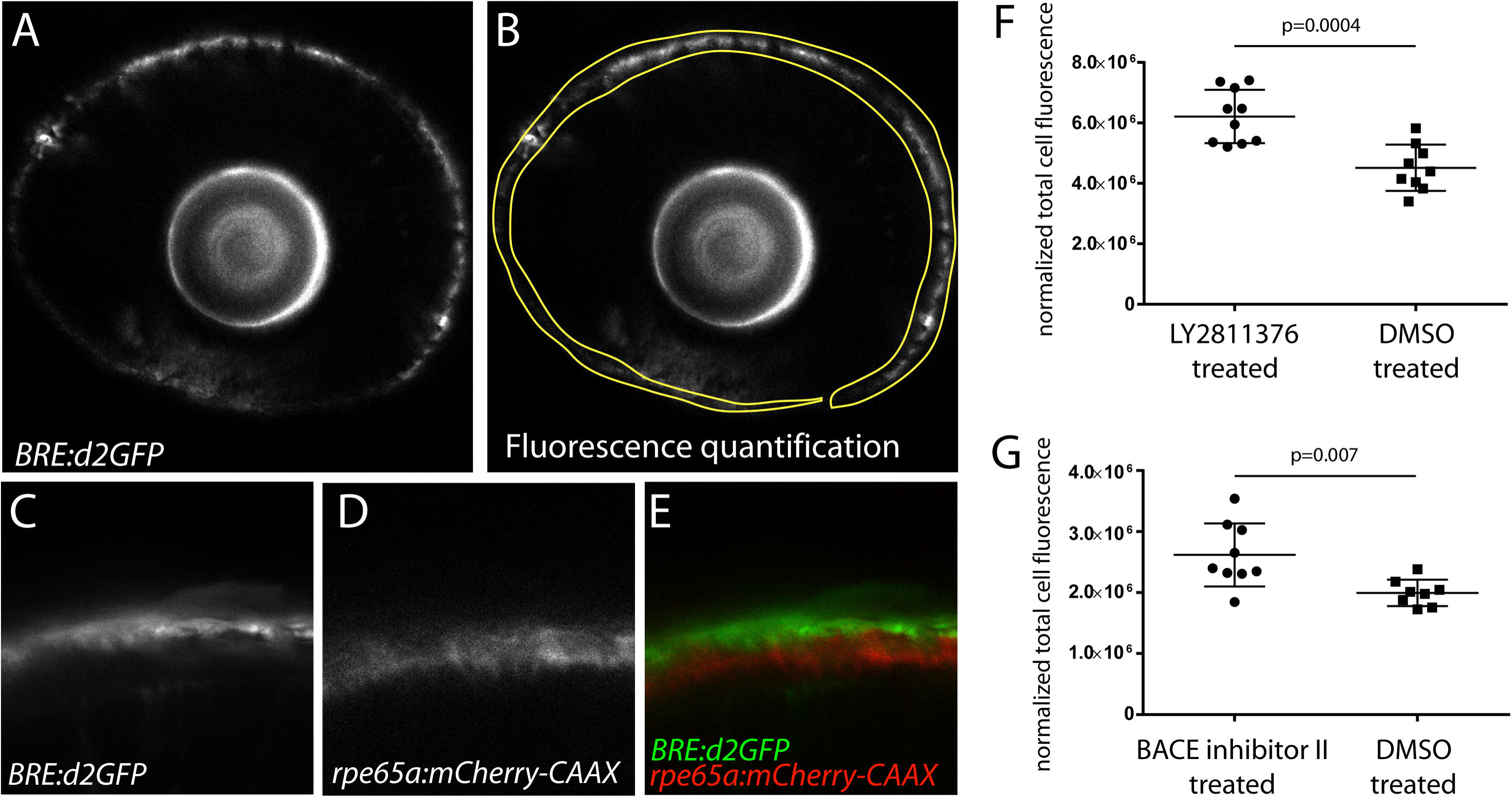
Inhibition of BACE enzyme activity in zebrafish increases Bmp signaling in the choroid/sclera of the eye. Incubation of 4-5 dpf *BRE:d2GFP* larvae with LY2811376 or BACE inhibitor II caused increased levels of Bmp signaling in the choroid/sclera of the eye compared to DMSO-treated control siblings, n = 10 for LY2811376 treatment; n= 9 for BACE inhibitor II treatment; n = 9 for DMSO control treatment.

To clarify the specific identity of the outer layers of the eye demonstrating augmented Bmp signaling in response to BACE inhibition, we incrossed *BRE:d2GFP* and *rpe65a:mCherry*-*CAAX* zebrafish and examined doubly transgenic larvae at 4 dpf. Green fluorescent tissue layers were found to lie outside the red fluorescent marked RPE layer (Figure 7 C - E), indicating that the sclera/choroid layers are responsible for the altered BMP signaling following inhibition of BACE enzyme activity.

### Expression of soluble, but not membrane-bound, Lrp2 LDLA1 domain from the RPE reduces choroidal/scleral Bmp activity

In order to investigate whether cleavage of Lrp2 from membrane-bound to soluble form affected its ability to modulate Bmp signaling, we made plasmids *rpe65a:lrp2 LDLA1*-*eGFP* and *rpe65a:lrp2 LDLA1*-*ICD*-*eGFP* that drive expression of either soluble Lrp2 LDLA1 or membrane-bound Lrp2 LDLA1 with its native transmembrane domain, respectively. These plasmids were separately injected into BRE:dmKO2 reporter zebrafish, and Bmp activity levels measured in the eye at 5 dpf by fluorescence quantification. In BRE reporter fish, fluorescence is almost completely confined to the choroid/scleral layers (see previous section). When compared with uninjected control *BRE:dmKO2* eyes, soluble Lrp2 LDLA1 reduces Bmp activity significantly, while membrane-bound Lrp2 LDLA1 has did not affect reporter activity (Figure 8). This indicates that soluble LDLA1 can act as an inhibitor for Bmps, preventing them from binding and signaling through appropriate receptors, while membrane-bound LDLA1 does not interfere with Bmp signaling, but potentially augments other pathways within RPE cells‥

**Figure 8.**
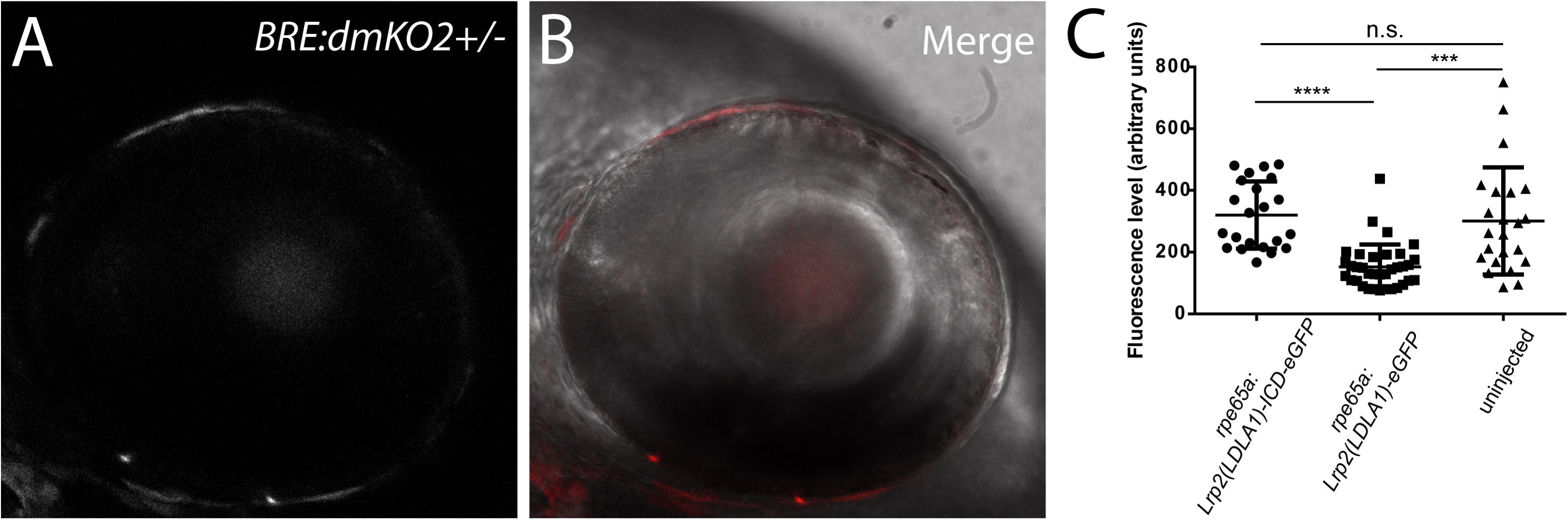
Expression of soluble, but not membrane-bound, Lrp2 LDLA1 from RPE downregulates Bmp signaling in the choroid/sclera. A, B *rpe65a:Lrp2 LDLA1*-*eGFP* or *rpe65a:Lrp2 LDLA1*-*ICD*-*eGFP* plasmids were injected into BRE:dmKO2 reporter zebrafish and eye fluorescence measured at 4 dpf. C. Sclera/choroidal expression levels were unchanged from uninjected controls with Lrp2 LDLA1-ICD-eGFP (uninjected, 301 arbitrary units ± 173; Lrp2 LDLA1-ICD-eGFP, 320 arbitrary units ± 109), while Lrp2 LDLA1-eGFP expression led to significant reduction of Bmp activity (153 arbitrary units ± 73). Fluorescence levels were compared using one-way ANOVA with Tukey’s post-hoc test.

### *lrp2*^-/-^zebrafish eyes have elevated transcription of ECM components and modulating genes

Associations between refractive error and genes involved in ECM growth and remodeling have been reported in individual gene studies, and in genome-wide association studies (25,26,58–64). In the eye, BMPs have been shown to inhibit excessive TGF-β2 induction of ECM proteins (65–69). Since we have shown that absence of Lrp2 leads to modulation of genes influenced by Bmp overexpression, we hypothesized that Lrp2 may assist in fine-tuning Bmp levels to properly control deposition and turnover of ECM proteins. GSEA analysis of *lrp2*^-/-^ and wild-type eyes at 1 mpf showed upregulation of ECM components and modulators when queried using ECM-receptor interaction pathway signature (KEGG pathway hsa04512; gene names converted from human to zebrafish) (Figure 9). This suggests that Lrp2 is necessary in the eye to maintain appropriate levels of ECM proteins in the eye, and that ECM protein dysregulation may result in the excessive eye growth seen in *lrp2/bugeye* mutants.

**Figure 9.**
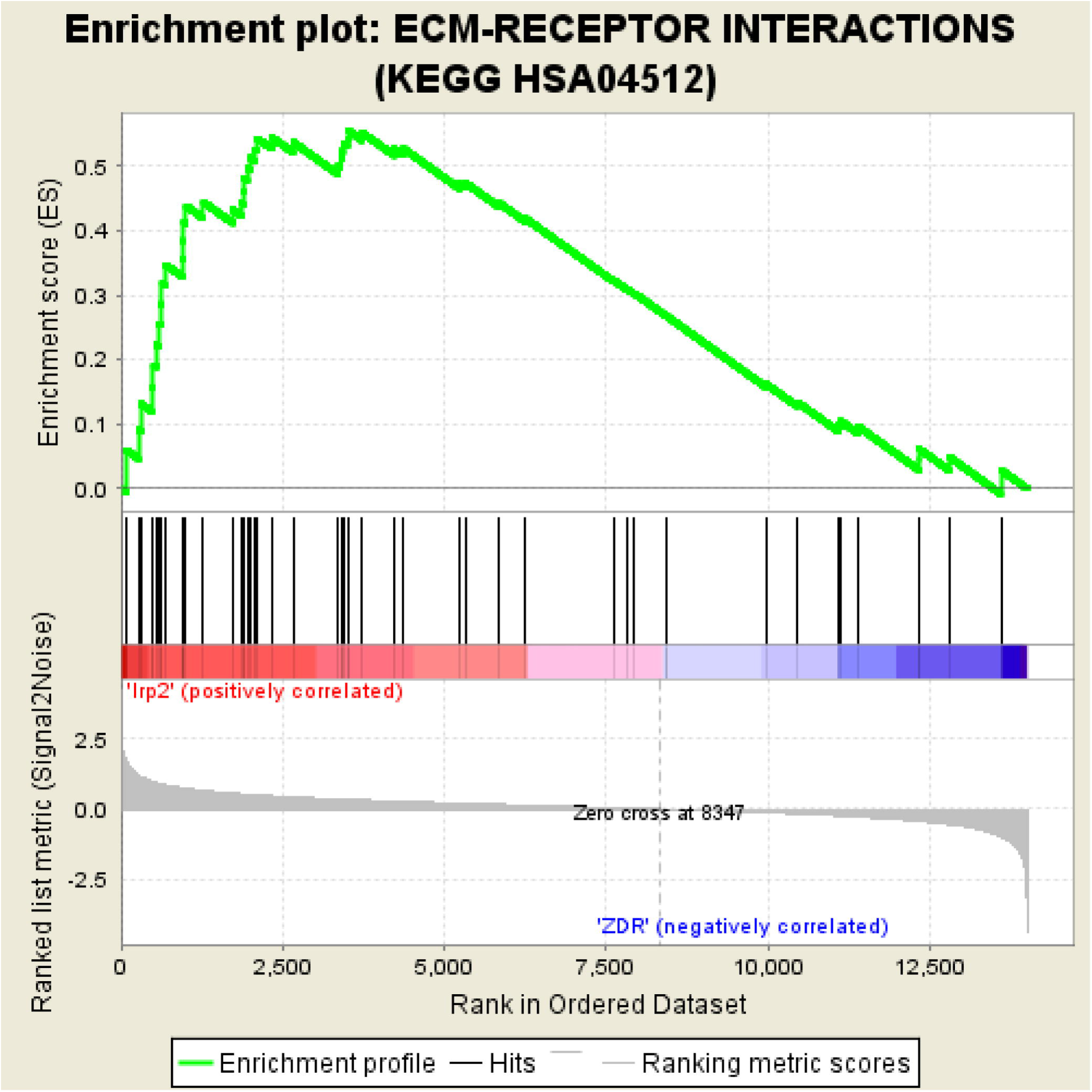
Expression of ECM components is upregulated in *lrp2*^-/-^ eyes. GSEA analysis using an ECM-receptor KEGG pathway converted to zebrafish orthologs to query 1 mpf eyes show enrichment of ECM gene expression with a normalized enrichment score = 1.897 for *lrp2*^-/-^ eyes (nominal p value = 0.002, FDR q = 0.002, FWER p = 0.001).

## Discussion

Mutations in Lrp2 cause excessive eye growth and myopia; however, the nature of the influence of Lrp2 on ocular globe size and emmetropization is likely multifactorial and has not been fully elucidated. Here we show that transgenic expression of recombinant, full length Lrp2 from the zebrafish RPE can significantly reduce the *lrp2* mutant phenotype, demonstrating that the effects of Lrp2 are at least partly local to the eye, and not exclusively the result of systemic factors that circulate in the blood or lymphatic systems. This interpretation is consistent with recent studies of mice with *foxg1*-mediated inactivation of Lrp2, which eliminates expression from the neural retina, RPE, and forebrain and caused eye enlargement and myopia (70). Using transcriptomic analyses, we found deletion of Lrp2 altered Bmp signaling, as well as other pathways. Because of the specificity of perturbed Bmp signaling to the RPE and sclera/choroid, coupled with previous studies implicating BMP signaling in emmetropization, we focused subsequent experiments on how Lrp2 regulates this pathway. Our studies indicate that Lrp2 can be cleaved from the RPE to release an extracellular domain capable of binding and inhibiting Bmp4 activity, and potentially activity mediated by other Bmps. Indeed, RPE expression of the Lrp2 LDLA1 domain mimicked the effects of known BMP2/4/7 ligand trap antagonists to promote eye size and myopia. Application of sheddase inhibitors to zebrafish larvae suggests that the Lrp2 extracellular domain is released from the basal side of the RPE, adjacent to the sclera /choroid, and cleavage is mediated by BACE activity. The Lrp2 sheddase, therefore, acts as a molecular switch to convert the full-length Lrp2 protein from a potential facilitator of Bmp signaling and modulator of other pathways, to a soluble antagonist of Bmp signaling. Cumulatively, this suggests that regulated cleavage of Lrp2 enables fine-tuning of multiple pathways to control ocular growth and emmetropization. We propose a model where wild-type eyes balance cleavage of Lrp2 from a membrane-bound to a soluble form thus fine-tuning Bmp signaling. When Lrp2 is completely absent, signaling pathways augmented by membrane-bound Lrp2, such as BMP, Shh, and perhaps others, can no longer be facilitated, leading to reduced activity and excessive eye growth (Figure 10). This disease phenotype can be further exacerbated by transgenic expression of soluble Lrp2 LDLA1 which acts as a decoy for Bmp ligands, decreasing signaling further and causing greater eye growth.

**Figure 10.**
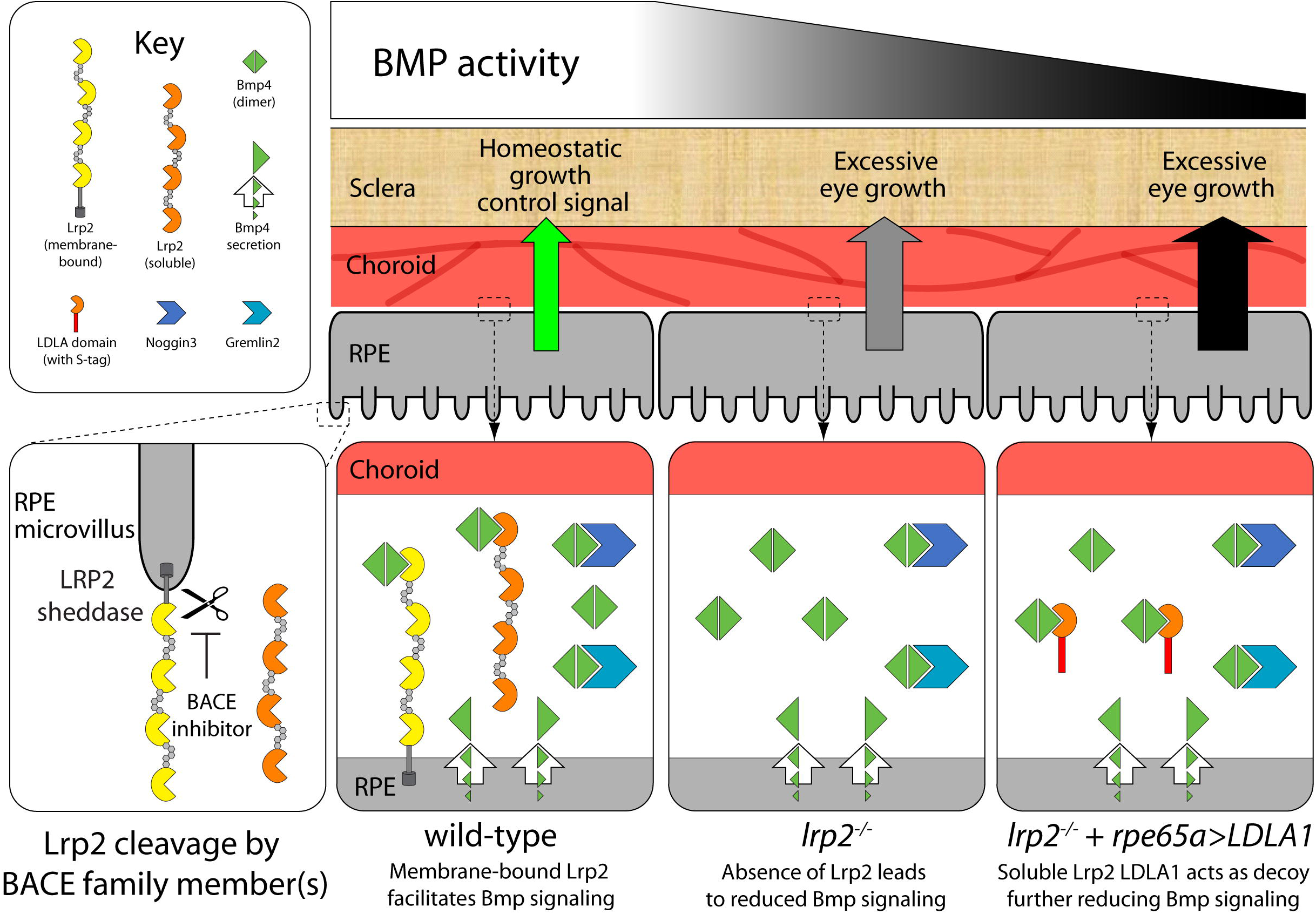
Model – Modulation of Lrp2-mediated Bmp4 signaling by extracellular cleavage allows fine-tuning of ligand levels between the RPE and sclera/choroid. Lrp2 may facilitate Bmp4 signaling while the extracellular region of Lrp2 is tethered by its transmembrane domain. Extracellular cleavage of Lrp2 by member of the BACE protein family converts the soluble domain to a decoy receptor that no longer facilitates signaling. Absence of membrane-bound Lrp2 leads to reduced Bmp signaling which is associated with excessive eye growth. When *lrp2*^-/-^ eyes with already reduced Bmp signaling express transgenic Lrp2 LDLA1 from the RPE, some Bmp ligands are bound by this decoy, further reducing Bmp signaling and increasing excessive eye growth even more.

Complexity in Lrp2 function is underscored by its effect on multiple pathways, but also on its distinct actions in different tissues. For example, in addition to altered Bmp signaling, loss of Lrp2 in zebrafish also affects Shh signaling in the eye. Our observed changes in Shh signaling with zebrafish are consistent with recent analysis of the retinal margin in *Lrp2* mutant mice (9). This study showed elevated Shh activity and subsequent hyperplasia of the ciliary epithelium. Expansion of the ciliary epithelium is also observed in zebrafish (43). In both mice and zebrafish with *lrp2* mutations, neural retinal growth, however, is insufficient for the size of the eye enlargement and the neuronal layers become stretched and thin prior to detected degeneration of ganglion cells. These observations suggest that Shh contributes to buphthalmia by promoting proliferation of the ciliary epithelium, which consequently increases aqueous humor production, and that fluid pressure promotes expansion of the ocular globe. Distinct from Lrp2 function in the ciliary epithelium, our data indicates RPE-localized Lrp2 modulates signaling that ultimately controls extracellular remodeling associated with the sclera and choroid which is known to underlie axial length changes.

Potentially, tissue-specific functions of Lrp2 may depend on the presence of ligands and expression of other cell surface (co-)receptors, as well as sheddase activity. As previously demonstrated, cellular and tissue context governs whether Lrp2 positively or negatively impacts Shh signaling (9,10,56). Lrp2 in the forebrain serves as a Patched co-receptor to augment Shh activity, while in the ciliary epithelium Lrp2 mediates Shh endocytosis and lysosomal shuttling to temper Shh activity. Similar complexity is emerging with the influence of Lrp2 on Bmp signaling. Previously, it has been demonstrated in forebrain ventricular cells that full-length Lrp2 can also endocytose and facilitate lysosomal degradation of Bmp4 (56). Conversely, full-length Lrp2 on RPE cells may function to augment Bmp activity within those cells.

In vascular endothelial cell precursors, the highly homologous Lrp1 can function as a co-receptor with Alk6/BmpRII for Bmp4 to promote holocomplex endocytosis to prolong and enhance Bmp signaling (71). This possibility is consistent with a requirement of Bmp activity implicated by myopia-associated variants that affect genes encoding ligands, receptors and regulators of Bmp signaling. Our studies indicate that cleavage of Lrp2 would disrupt full-length function(s) and convert the extracellular domain to a Bmp ligand trap antagonist. In this manner, then, the fraction of uncleaved Lrp2 on RPE cells could autonomously augment Bmp signaling, while shed extracellular protein would non-autonomously temper Bmp signaling in the sclera and choroid. Proteolytic processing would also affect Shh signaling by preventing the full-length functional influence of Lrp2 on that pathway.

Our discovery that the shed extracellular domain of Lrp2 is functional has several implications. First, modulation of the BACE(-like) sheddase might serve as a therapeutic target for pathological, progressive myopia. However, the BACE1 inhibitor LY2811376 has been shown to cause retinal dystrophy in rodents (72,73). Similarly, genetic deletion of BACE1 in mice resulted in significant choroidal, RPE, and retinal pathology (73). As BACE proteases are known to process many different proteins, any therapeutic approach to modulate Lrp2 cleavage would need precision. Related to cleavage of the Lrp2 extracellular domain, it is interesting to note that mouse models of Lrp2 knockout show different phenotypes depending on the mutant allele: homologous recombination to replace 1.5 kb of the *Lrp2* gene with a neomycin cassette results in near-total embryonic lethality in homozygous *Lrp2* mice due to holoprosencephaly, mimicking loss of Shh function. In contrast, a single point mutation causing a premature stop codon after amino acid 2721 allows for significantly greater survival and leads to eye enlargement and myopia (74–77). Prematurely terminating Lrp2 before the transmembrane domain may produce exclusively the soluble extracellular domain, restricting Lrp2 activity to a ligand trap function. Specifically engineered mutant alleles for Lrp2 should help illuminate the multi-modal activities of Lrp2.

Importantly, we show that transcriptional upregulation of ECM components in the eye is one outcome of *lrp2* inactivity, which directly suggests a mechanism for excessive eye growth and refractive error to occur. Overall, our work along with that of others supports the concept that Lrp2 is a homeostatic node that buffers and integrates multiple pathways to precisely control initial eye growth and emmetropization. In its absence, subtle and sustained alterations in signaling impact eye size and refraction and cause severe myopia.

## Materials and Methods

### Fish maintenance

Zebrafish (*Danio rerio*) were maintained at 28.5°C on an Aquatic Habitats recirculating filtered water system (Aquatic Habitats, Apopka, FL) in reverse-osmosis purified water supplemented with Instant Ocean salts (60 mg/l) on a 14 h light: 10 h dark lighting cycle and fed a standard diet (78). All animal husbandry and experiments were approved and conducted in accordance with the guidelines set forth by the Institutional Animal Care and Use Committee of the Medical College of Wisconsin.

### Transgenesis

DNA constructs for transgenesis were generated by PCR amplifying genes or domains of interest from adult eye cDNA (*nog3* CDS, *grem2* CDS, *bmp4* CDS, full-length and domains of *lrp2*) or gDNA (*rpe65a* promoter) before using the Invitrogen Gateway system (Thermo Fisher Scientific, Waltham, MA) to generate entry clones. The Gateway system was further used to assemble promoter: gene-tag constructs using backbone vectors containing Tol2-inverted repeats flanking the transgene constructs to facilitate transgenic insertion into the zebrafish genome (79,80). Plasmid and primer sequences are available on request.

### Spectral domain-optical coherence tomography (SD-OCT)

Zebrafish eyes were imaged using a Bioptigen Envisu R2200 SD-OCT imaging system with a 12 mm telecentric lens (Bioptigen, Morrisville, NC) essentially as described (46). To ensure uniform measuring of eyes, the iris edges were aligned in both dorsal-ventral and nasal-temporal planes before acquisition. Ocular dimensions were measured using the built-in manual caliper tool in the InVivoVue software platform with an applied refractive index constant of 1.30.

### RNA extraction and RNAseq

*lrp2*^-/-^ and wild-type zebrafish eyes were dissected at 1 mpf to give retinal, RPE and sclera/choroid samples. Each sample contained tissues from 10 – 20 eyes and was repeated in triplicate. RNA extracted from these samples was assayed for tissue specificity using retinal-, RPE- or sclera/choroid-specific marker genes by qPCR, before analyzing by RNAseq. RNA was processed for sequencing by BGI Americas (Cambridge, MA), using an Illumina HiSeqTM 2000 instrument, and Bowtie2 used to align to the current zebrafish genome (GRCz10 (GCA_000002035.3)).

### Direct link to deposited data

The datasets supporting this article are available in the NCBI Gene Expression Omnibus (GEO) archive. Accession number for the data is GSE97125. Link to the data: https://www.ncbi.nlm.nih.gov/geo/query/acc.cgi?acc=GSE97125.

### Gene set enrichment analysis (GSEA)

RNAseq datasets were analyzed for enrichment of genes associated with transcript signatures defined by overexpression of zebrafish Bmp4 and Shha. Briefly, datasets and signature gene lists were entered into the stand-alone GSEA tool (48,49). Resulting sets of enriched genes were tabulated using standard output criteria (enrichment score (ES), normalized enrichment score (NES), nominal p-value, false discovery rate q-value, family-wise error rate).

### HEK293T cell culture and immunoprecipitation

Zebrafish Lrp2 LDLA domains 1 through 4 were individually placed under the control of a CMV promoter and tagged at their C-terminus with a 15 amino acid S•tag epitope, which can be immobilized using S•protein conjugated to agarose beads. The native Lrp2 signal peptide was placed upstream of each LDLA domain to facilitate proper protein export. Zebrafish Bmp4 sequence was placed downstream of a CMV promoter, and an internal eGFP tag was placed after the furin cleavage sites so that when translated mature Bmp4 could be detected after removal of the prodomain (81). Plasmids *CMV:eGFP*-*bmp4* and *CMV:lrp2 (signal peptide*-*LDLAx)*-*S*•*tag* (where x represents one of the LDLA domains 1 – 4) were transfected into HEK293T cells for 24 hours. Proteins were extracted from pelleted cells using a gentle RIPA buffer, and incubated with S-protein agarose (EMD Millipore, Temecula, CA) before washing and eluting from agarose by boiling in reducing protein loading dye. Proteins were separated on Bio-Rad Any kD gradient SDS-PAGE gels before transferring to Immobilon-F PVDF membranes. Western blots were probed using anti-eGFP antibody JL-8 (Clontech, Mountain View, CA) and developed using the LI-COR Odyssey buffer and imaging system (LI-COR, Lincoln, NE).

Similarly, HEK293T cells were transfected with *pTol2*-*CMV:SP6*-*seGFP*-*lrp2 ECD*-*ICD*-*mCherry* plasmid DNA and cultured for 24 hours before adding inhibitors or activators of sheddases. These compounds were incubated with the cells for 24 hours before conditioned medium was collected. Conditioned medium was applied to GFP-Trap^®^_A agarose (Chromotek, Hauppauge, NY) for 2 hours before washing the agarose and eluting bound protein by boiling in reducing protein loading dye. Western blotting was carried out as above. Where appropriate, Western blot bands were quantified using ImageJ (Rasband, W.S., ImageJ, U. S. National Institutes of Health, MD).

### Zebrafish in vivo Bmp reporter assay

BACE inhibitor compounds or DMSO as control were bath-applied to transgenic *BRE:d2GFP* zebrafish for 24 hours before imaging fluorescence in the eye using a Nikon Eclipse E600FN confocal fluorescent microscope (Nikon, Tokyo, Japan). Fluorescent intensity was measured in ImageJ by outlining the region of interest and measuring the integrated density for fluorescence, correcting for background fluorescence (82). Double transgenic *BRE:d2GFP/ rpe65a:mCherry*-*CAAX* or *BRE:dmKO2* zebrafish were imaged at 5 dpf to identify the GFP-positive tissue in the larval eye.

### Statistical analyses

Eye measurements were processed using Microsoft Excel (Microsoft, Redmond, WA) and graphed using GraphPad Prism (GraphPad, La Jolla, CA). Standard deviation (SD) and analysis of variance (ANOVA) with post-test analyses were calculated using GraphPad Prism. Sample sizes were not calculated in advance, but post hoc calculations showed that most experiments exceeded a statistical power of 95% for alpha = 0.05. Post hoc sample sizes were calculated for axial length, axial length: body axis, and axial length: lens diameter metrics, to detect an outcome of 10% change relative to wild-type at 2 mpf and 3 mpf. An exception was 3 mpf wild-type axial length: lens diameter, where a 20% size change could be detected with significance using the numbers of animals tested. *lrp2*^-/-^ mutant eye metrics usually changed by more than 10% of the wild-type metric under consideration.

## Acknowledgements

The authors thank Michael Cliff and William Hudzinski for zebrafish husbandry. We are also grateful to Amy Ludwig-Kubinkski, Jason Bader and Amira Pavlovich for assistance and advice on cell culture and Western blotting. Finally, we are indebted to Dr. Joe Besharse for critical feedback and suggestions. This work was supported by National Institutes of Health/National Eye Institute R01 research grant EY016060 (BAL) as well as a Core Grant for Vision Research (P30 EY001931).

## Competing Interests

The authors have no competing interests of any kind to disclose.

## Figure Legend

**SI Figure. qPCR with tissue-specific markers shows high levels of purity for dissected sclera/choroid, RPE and retina.** Dissected sclera, RPE and retina were assayed for expression of genes enriched in those tissues (*col1a1a*, *col12a1a*, *sclera*/*choroid*; *rpe65a*, *rlbp1b*, RPE; *gnat2*, *pde6c*, retina). Enrichment levels for each tissue are shown as normalized ΔΔCq (A), or as percentages (B).

**S1 Table. RNAseq datasets comparing wild-type and *lrp2*^-/-^ zebrafish whole eyes, choroid/sclera, RPE and retina at 1 mpf, filtered for p<0.01, 50 RNAseq reads, and representation in all three sample replicates.**

**S2 Table. FPKM (Fragments Per Kilobase Million) gene transcript levels for 1 mpf wild-type and *lrp2*^-/-^ zebrafish eyes and dissected eye tissues.**

**S3 Table. RNAseq datasets comparing wild-type and *lrp2*^-;-^ zebrafish whole eyes, choroid/sclera, RPE and retina at 1 mpf, filtered for p<0.01, 50 fragments per kilobase of transcript per million mapped reads, and representation in all three sample replicates analyzed with Bmp4 and Shha signature pathway genes from this work.**

**S2 Figure. Bmp4 gene signatures used in this study.** A, B. Genes up- or down-regulated in 5 dpf eyes from *hsp70l:eGFP*-*bmp4* transgenic larvae compared with non-transgenic sibling controls were filtered for representation in three independent samples, and used to define zebrafish Bmp4-UP and Bmp4-DOWN gene signatures. C, D. Gene lists of the signature genes identified in A and B.

**S3 Figure. Shha gene signatures used in this study.** A, B. Genes up- or down-regulated in 5 dpf eyes from *hsp70l:shha*-*eGFP* transgenic larvae compared with non-transgenic sibling controls were filtered for representation in two independent samples, and used to define zebrafish Shha-UP and Shha-DOWN gene signatures. C, D. Gene lists of the signature genes identified in A and B.

**S4 Figure. Internal tagging of Bmp4 with eGFP allows *in vivo* tracking of the morphogen without loss of activity.** A. Schematic of Bmp4 showing insertion of eGFP downstream of furin cleavage sites tagging the mature ligand. B. Multiple cleavage forms of eGFP-Bmp4 can be seen on Western blots of larval zebrafish extracts expressing eGFP-Bmp4 under the control of a heat-shock inducible promoter. The bands likely correspond to full-length propeptide eGFP-Bmp4, mature cleaved eGFP-Bmp4, and intermediates. C - F. eGFP-Bmp4 is secreted from cells in the developing embryos following expression. G, H. Overexpression of eGFP-Bmp in the developing embryos causes a ventralized phenotype. I –T. Overexpression of eGFP-Bmp4 in BRE-dmKO2 show perdurance of secreted Bmp4 24 hours after transgenic expression.

**S5 Figure. *rpe65a*-driven eGFP-Bmp4 causes larval coloboma and severe eye disorganization in adult zebrafish.** A, E. rpe65a-derived eGFP-Bmp4 causes larval phenotypes including coloboma. The failure of the optic fissure to close properly is often accompanied by prognathia of the mandible. Most larvae die, but some survive to adulthood, and show severe eye disorganization. F – L. Coloboma is seen where the optic fissure remains open with the lens protruding. Staining with anti-cone arrestin 3 antibody, zpr-1, shows normally laminated cones in wild-type adults, but disorganization in plasmid-injected adults, where photoreceptor rosetting is seen.

**S6 Figure. RNAseq shows enriched expression of Bmp ligands in the RPE and choroid/sclera.** Bmp ligand family members, Bmp receptor family members and activin family members were assayed for expression levels in whole eyes and dissected eye tissues.

**S7 Figure. Zebrafish Lrp2 contains putative consensus substrate sites for BACE enzyme cleavage.** CLUSTAL alignments between zebrafish Lrp2 and published substrate sites of BACE show some homology. Locations of putative cleavage sites are indicated on a schematic representation of Lrp2.

